# JMJD6 and YBX1 physically interact and regulate HOTAIR proximal promoter

**DOI:** 10.1101/2024.12.28.630568

**Authors:** Aritra Gupta, Siddharth Bhardwaj, Kartiki V. Desai

## Abstract

In a previous study, we showed JMJD6 interacted with HOTAIR promoter (−123 to -103 bp upstream of TSS, JIR) and augmented its transcription. Maximal JMJD6 mediated induction required (−216 to -123 bp) region. *In silico* prediction and ENCODE data suggested that YBX1 could be that potential candidate and in this study the region is designated as YBX1 interacting region (YIR). In breast cancer cell lines, we show that JMJD6 and YBX1 regulate each other’s expression and physically interact with each other when recombinantly expressed, as endogenous proteins and when synthesized *in vitro.* Domain mapping indicated that A/P domain of YBX1 interacted with JMJC domain of JMJD6. Luciferase activity of HOTAIR promoter constructs, pHP216 and pHP123, increased in presence of YBX1 in MCF7, Vec and JMJD6 overexpressing JOE cells but was lost in the presence of JMJD6 and YBX1 siRNAs. Interestingly, activity of pHP123 that lacks YIR also decreased upon YBX1 knock out (YKO). Next, by individual JMJD6, YBX1 and ChIP-re-ChIP assays we demonstrate that both proteins co-occupy this promoter region. Further, electrophoretic mobility shift assays showed that YIR probes retarded two complexes, which lost intensity in YKO cells. Interestingly, JIR-protein complex disappeared in YKO cells. Together these data imply that YBX1 not only enhanced promoter activity but may also be involved in JMJD6 recruitment. Taken together, our data proposes that the interaction and positive feed forward loop perpetuated by JMJD6 and YBX1 may culminate in HOTAIR induction, which in turn is known to drive tumor progression.

## Introduction

Jumonji domain containing protein 6 (JMJD6) is an arginine demethylase and lysyl hydroxylase that epigenetically regulates gene transcription (1–4). We have earlier identified it as a poor prognostic factor in breast cancer and its positive impact on cell proliferation and motility (5). In addition, we showed that JMJD6 and the long intervening non-coding RNA (lincRNA) HOTAIR are co-expressed in breast tumor samples and their combined high expression promotes poorer survival in patients (6). In the same study, a JMJD6 interaction region (JIR), that positively regulated HOTAIR expression in breast cancer cell lines was identified. JIR was found approximately -123 to -103 bases upstream of the HOTAIR transcription start site (TSS). Addition of a further upstream 100bp region (−216 to -123 bp) increased this basal promoter activity in JMJD6 overexpressing cells (JOE). Several potential transcription factor sites were predicted across this region and mutations of potential sites failed to identify the nature of interacting protein(s). However, these regions were positive for gel-shifts using nuclear extracts from JOE cells suggesting the presence of other regulatory factor(s).

Though JMJD6 has a RNA binding activity, it lacks a DNA binding domain. However, it interacts with several transcription factors including bromodomain containing protein 4 (BRD4) and MYC, and has emerged as a key player in anti-pause release of stalled RNA pol II sites to augment gene transcription (7). While JMJD6 hydroxylates p53 negatively affecting its target gene transcription, it improves translocation of ER from the cytoplasm to nucleus by virtue of its arginine demethylase activity (8,9). Further, JMJD6 is recruited to ER binding sites and is required for estrogen induced gene transcription in breast cancer cells (10). We recently showed that JMJD6 decreased ER expression and promoted insensitivity to endocrine therapy drugs such as Tamoxifen (11). Here, JMJD6 hijacked the E2-ER axis of proliferation by inducing E2F regulated genes and G2-M transitions to sustain breast cancer growth despite loss of ER expression. Considering the versatility of JMJD6 action, it is possible that it interacts with additional transcription factors to mediate gene expression.

In this manuscript, we combined ENCODE data and molecular studies to identify Y-box binding protein 1 (YBX1) as a possible factor that interacts with this region of the basal HOTAIR promoter (12). We reviewed the mass spectrometry data obtained using anti-JMJD6 antibodies by Webby et al and Liu et al, and found YBX1 as one of the potential interactors in the list of immunoprecipitated proteins (4,7). Based on its functional similarity with JMJD6, such as association with poor prognosis, participation in mRNA splicing via U2AF factors, its ability to bind RNA and decrease ER expression and it’s DNA binding activity as CCAATT-enhancer binding protein, we envisage that JMJD6 and YBX1 co-regulate cancer related genes(13–17). Previous reports also show that HOTAIR in turn augments nuclear translocation of YBX1 and its oncogenic activity raising the possibility of a regulatory loop in these proteins (18). Here, we show that JMJD6 and YBX1 are expressed in a positive feed forward loop, both proteins interact with each other physically and confirm that YBX1 interacts with the HOTAIR promoter to enhance its expression.

## Materials and Methods

### Cell Culture

MCF7, MDA MB 231, HEK293 cells were purchased from American Type Culture Collection (ATCC, VA, USA). The cell lines were grown in Dulbecco’s Modified Eagle’s Medium (DMEM) (GIBCO, USA) with 5% fetal bovine serum (GIBCO, USA) and 1% Penicillin-streptomycin (GIBCO, USA) in humidified 5% CO2 incubator at 37^0^C. MCF7 cells stably expressing recombinant JMJD6 (referred as JOE cells) and control MCF7 cells carrying only the empty vector (Vec) have been described by us earlier (5). Stable clones were maintained in 750µg/ml Geneticin (Thermo Scientific, USA).

### Over expression of JMJD6 and YBX1

Construction of JMJD6-V5 plasmid was previously summarized and pDEST Myc (#19878) tagged YBX1 plasmid was purchased from Addgene (5). PCR based deletion constructs of JMJD6 and YBX1 were made using primers described in Supplementary table 1. Plasmids were transiently (co)-transfected in various cell lines using standard protocols and Lipofectamine 2000 (Thermo Scientific, USA) for various experiments detailed below.

### Depletion of JMJD6 and YBX1

SiRNA mediated knock down (KD) of JMJD6 and YBX1 was performed using reverse transfection method. JMJD6 siRNA (5’GCUAUGGUGAACACCCUAATT 3’), YBX1 siRNA (5’ GTTCAATGTAAGGAACGGAT3’) (5,19) were synthesized (Eurogentec) and for control non-targeting control (si_Scrambled RNA) was used (Ambion). CRISPR based knock out (KO) of JMJD6 and YBX1 was conducted. sgRNA with minimal off target effect was selected using CRISPOR tool and cloned in e-sp-Cas9-2a-GFP vector. Pooled KO clones were used for experiments since KO of JMJD6 and/or YBX1 proved to be lethal for cells in our hands.

### Western Blot

Cells were lysed in RIPA buffer, with 1x Protease inhibitor cocktail (PI) (Sigma Aldrich) and protein extracts were quantified by the BCA method (Pierce, USA). Equal amounts (50 micrograms) of proteins were analyzed in 10-15% SDS-PAGE gels, transferred on to PVDF membrane (Millipore, Germany) and nonspecific sites were blocked by using 5% non-fat milk (Bio-Rad, USA). Primary antibodies were incubated at 4° C overnight and after washing, blots were probed with suitable HRP-conjugated secondary antibody for 1 hour at room temperature. Signals were detected using HI-ECL (Bio-Rad, USA) in Chemidoc (Bio-Rad, USA). Antibodies used for Western blots were JMJD6 for total protein (endogenous and exogenous) (PSR-H7, Santacruz Biotechnology sc-28348, USA, 1:1000), V5 for exogenously expressed JMJD6-V5 (Thermo Scientific, USA, R960-25,1:5000), YBX1 (exogenous and endogenous) (Abcam, UK, ab-12148, 1:2000), Myc for exogenously expressed Myc-YBX1 (Sigma Aldrich, USA, 05-419, 1:2000), β-Actin as internal control (Thermo Scientific, USA, A2228, 1:5000), Lamin A/C (Abclonal, USA, 1:5000, A0249), and β tubulin (Abclonal, USA, 1:5000, AC008) for nuclear and cytoplasmic markers respectively. Densitometric scanning of 3 independent blots at variable exposures was conducted and data is expressed as average numeric values under western blots wherever applicable.

### Immunofluorescence microscopy

5×10×4 cells were plated onto poly-D-lysine coated coverslips for 24 hours. 10% neutral buffer formalin was used for fixation, after washing, cells were subjected to permeabilization (PBS + 0.2% Tween20) and a Glycine wash. Non-specific sites were blocked using 10% goat serum for 1 hour. Cells were incubated over night with primary antibody, washed and incubated with fluorophore tagged secondary antibody for 1.5 hours (Abcam, UK).

Coverslips were mounted using mounting media containing DAPI (Sigma Aldrich). Signals were captured in Confocal microscope (Nikon Eclipse, Ti2E). For proximity ligation assay (PLA) after primary antigen probing, Duo-link kit protocol was used (Sigma Aldrich). JMJD6 and YBX1 primary antibodies were used at suitable dilutions and PCR and secondary probing was done using reagents provided in the kit. Signals were captured at high resolution by Confocal microscope (Nikon Eclipse Ti2E).

### Cell fractionation

Cells were incubated with 5X volume of cytoplasmic extraction buffer supplemented with PI and DTT for 3 minutes for isolating cytoplasmic protein fraction. The lysate was centrifuged and the supernatant reserved as the cytoplasmic fraction. The pellet was washed with cytoplasmic extraction buffer without NP40. After measuring packed nuclear volume, nuclei were lysed in nuclear extraction buffer containing 1X PI and DTT for 10 minutes. All processes were conducted at 4°C.

### Promoter reporter assays

Previously reported constructs pHP216 and pHP123 containing (−216 to +50 bp) and (−123 to +50 bp) regions of HOTAIR promoter were used for assays in presence/absence of JMJD6/YBX1 or both. pRL-TK plasmid was used as a control for determining transfection efficiency. Cells were transfected with various constructs using Lipofectamine 3000 and harvested 48 hours post transfection. Dual Luciferase Assay kit (Promega Corporation, WI, USA) was used to estimate luciferase activity. Relative luciferase activity was calculated by comparing with Renilla luciferase activity.

### Co-immunoprecipitation (co-IP)

500 µg of proteins from freshly prepared protein lysates of relevant cell lines were precleared using protein A/G agarose beads (Santacruz, USA). After nutation of 4 hours, supernatant was mixed with antibody-bead slurry and kept overnight for nutation. Next day, after vigorous washing, proteins were eluted by boiling for 10 minutes in SDS-PAGE buffer in the presence of β-mercaptoethanol (BME) Immunoprecipitated complexes were detected by Western blots. 10% of extracts were reserved as input prior to co-IP. Veriblot (Abcam, UK) was used as a secondary antibody control.

### *In vitro* protein synthesis and interaction

JMJD6 and YBX1 constructs containing an upstream T7 promoter were linearized and used as a template for coupled transcription and translation in rabbit reticulocyte lysates according to manufacturer’s protocols (TnT kit,Promega Corporation, USA). Synthesis was confirmed using Western blot analysis using respective antibody*. In vitro* synthesized proteins were conjugated with respective antibody-A/G bead slurry. After 12 hours of nutation non-specific proteins were washed off, followed by overnight nutation with the partner protein. Immunoprecipitated complexes were detected as described above.

### Chromatin Immunoprecipitation (ChIP)

Cells were cross linked using formaldehyde solution and reaction was stopped using Glycine. For JMJD6 ChIP, Di succinimidyl gluterate (DSG) was additionally used as a crosslinker. For JMJD6 ChIP, V5 antibody and for YBX1, YBX1 antibody was used. Chromatin isolated was sheared to 300 to 700 bp fragments using a bath sonicator (Covaris S200). Fragment size was confirmed by D1000 screen tape on Agilent tape station. Antibody pull down was performed overnight with A/G magnetic beads, followed by standard washing and protein DNA de-crosslinking. For sequential ChIP, ChIP material from the first pull down was not de-crosslinked, instead, incubation with the partner protein antibody was continued. Isolated DNA was run through PCR purification columns before quantitative PCR with primers for selected chromatin regions (Supplementary table 1). 10% input DNA was purified prior to ChIP and used for enrichment analysis using the Percentage Input method.

### Electromobility shift assay (EMSA)

Nuclear extracts were freshly prepared from 10^7^ cells. Cyanine-5 Fluorescently labelled DNA probes (SIGMA chemical Co.) have been described earlier (Supplementary table 1) and the LUEGO protocol used for EMSA (20). 100 nM probes were incubated with 10 µg of nuclear extracts in a binding buffer (10 mM Tris–HCl, pH7.6, 50 mM KCL, 1 mM DTT, 10% glycerol, 200 ng/reaction poly(dI-dC)) for 30 min on ice. Reaction mixtures were loaded onto a non-denaturing 5% acrylamide: bisacrylamide (29:1) gel and run for 45 min at 100 V in 0.5× TBE buffer. 50× unlabelled cold competitor probe was used to determine specificity of complexes found. Gels were scanned after electrophoresis using ChemiDoc (Biorad) using the Cy-5 channel. Extracts from pooled CRISPR knock out clones were used as YKO protein samples.

### RNA isolation and quantitative RT-PCR (qRT-PCR)

RNA was isolated using RNAeasy isolation kit (Qiagen Corp.). 1 µg of RNA was taken for reverse transcription using random hexamer and Super script III (Thermo Scientific, USA). 1µl of cDNA was used to perform qRT-PCR in thermal cycler (Bio Rad) using specific primers (Supplementary table 1).

### Survival Analysis

Breast cancer data from KM plotter (www.kmplot.com/analysis) was utilized. For each gene, tumors expressing above median expression were grouped as high expressors and those below as low expressors. High expressors from all 3 genes were compared with tumors classified as low expressors for all 3 genes and probability of survival was estimated.

### Statistical analysis

All the experiments are done three times; graphs are made by Graph pad prism. Statistical analysis was done by R. Comparisons between two groups were performed using student’s t-test and data was considered statistically significant at p ≤ 0.05. Significance in all figures is indicated as follows: * p ≤ 0.05, **p ≤ 0.01, ***p ≤ 0.001 and ****p ≤ 0.0001.

## Results

### *In silico* analysis of HOTAIR proximal promoter

Our previous data indicated that JIR (−123 to -103 bp) and a separate region (−216 to -123 bp) of the HOTAIR promoter were required for its upregulation. The upstream site may contain binding site(s) for additional transcription factor(s) as determined by a gel-shift pattern showing potential interactors using DNA probes designed for (−216 to -123 bp) region (6).

Despite mutating predicted TF binding site(s) in this region, luc activity remained high and the factor was not identified. Here, we utilized *in silico* data from ENCODE having an experimental matrix of 160 TFs in MCF-7 cells. We found low enrichment profiles (1.5-2.5-fold) for 4 TFs; EP300, FOXA1, ZBTB11 and YBX1 in basal promoter region of HOTAIR. Of these, the former 3 factors could be eliminated, since our previous mutation study indicated that despite disrupting their cognate binding sites, the promoter activity remained unchanged. Analysis of published YBX1 ChIP data had yielded consensus binding sequences for multiple transcription factors (17). The same study reported that YBX1 may interact with sites like (5’-A/GCCATGG-3’), a YBX1 RNA consensus binding site identified in SAGE data. Such a partial site (5’-CATGG-3’) was found downstream of the site we had mutated earlier, and it overlapped with the YBX1 ENCODE data (Figure 1A). This site was explored further as a potential YBX1 interacting region (YIR).

**Figure 1:**
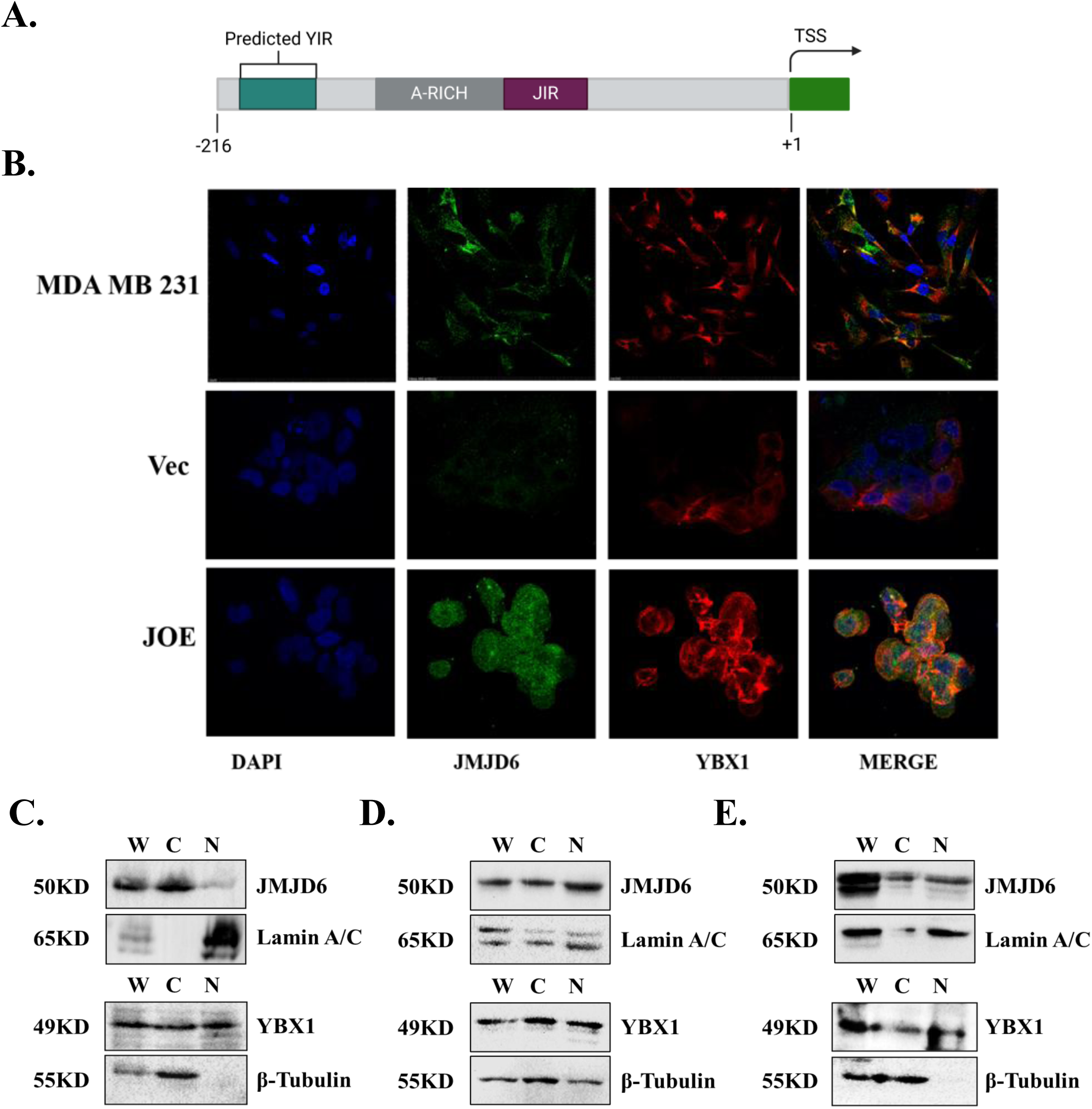
A) Schematic representation of predicted YBX1 interaction regions in HOTAIR promoter; B) Immunofluorescent detection of JMJD6 and YBX1 in MDA MB 231, Vec and JOE cells; C-D) Nuclear–cytoplasmic fractionation and immunoblots for JMJD6 and YBX1 in MDA MB 231 cells, and E-F) in JOE cells (W= whole cell lysate, C= Cytoplasmic extract, N= Nuclear extract).

### Expression levels of YBX1

Since the HOTAIR promoter was previously characterized in JOE, Vec and MDA MB 231 cell lines, YBX1 expression levels were determined by western blotting and immunofluorescence in these cells (Figure 1B, supplementary figure 1). Surprisingly, JMJD6 was found in the cytoplasm in MDA MB 231 and similar localization was shown in earlier reports (21,22). Abundant expression of YBX1 was found in all cell lines. Immunoblots were performed in lysates from nuclear and cytoplasmic fractions to localize these proteins.

JMJD6 showed prominent bands in the cytoplasmic fractions of MDA MB 231 but nuclear in Vec and JOE cells (Figure 1C, D and E respectively). On the other hand, YBX1 was found in both cellular compartments in all cells. Interestingly, YBX1 expression was elevated in JOE cells (Figure 1B, supplementary figure 1-right panel). Transient overexpression of JMJD6 in MCF7 cells also led to induction in level of YBX1 at both RNA and protein levels (Figure 2A-C). Conversely, overexpression of YBX1 elevated the level of JMJD6. SiRNA mediated knock down of either of the proteins reduced the expression of the other at both RNA and protein levels (Figure 2D-F). These data indicated that the two proteins may establish a positive feed forward loop by regulating each other’s expression.

**Figure 2:**
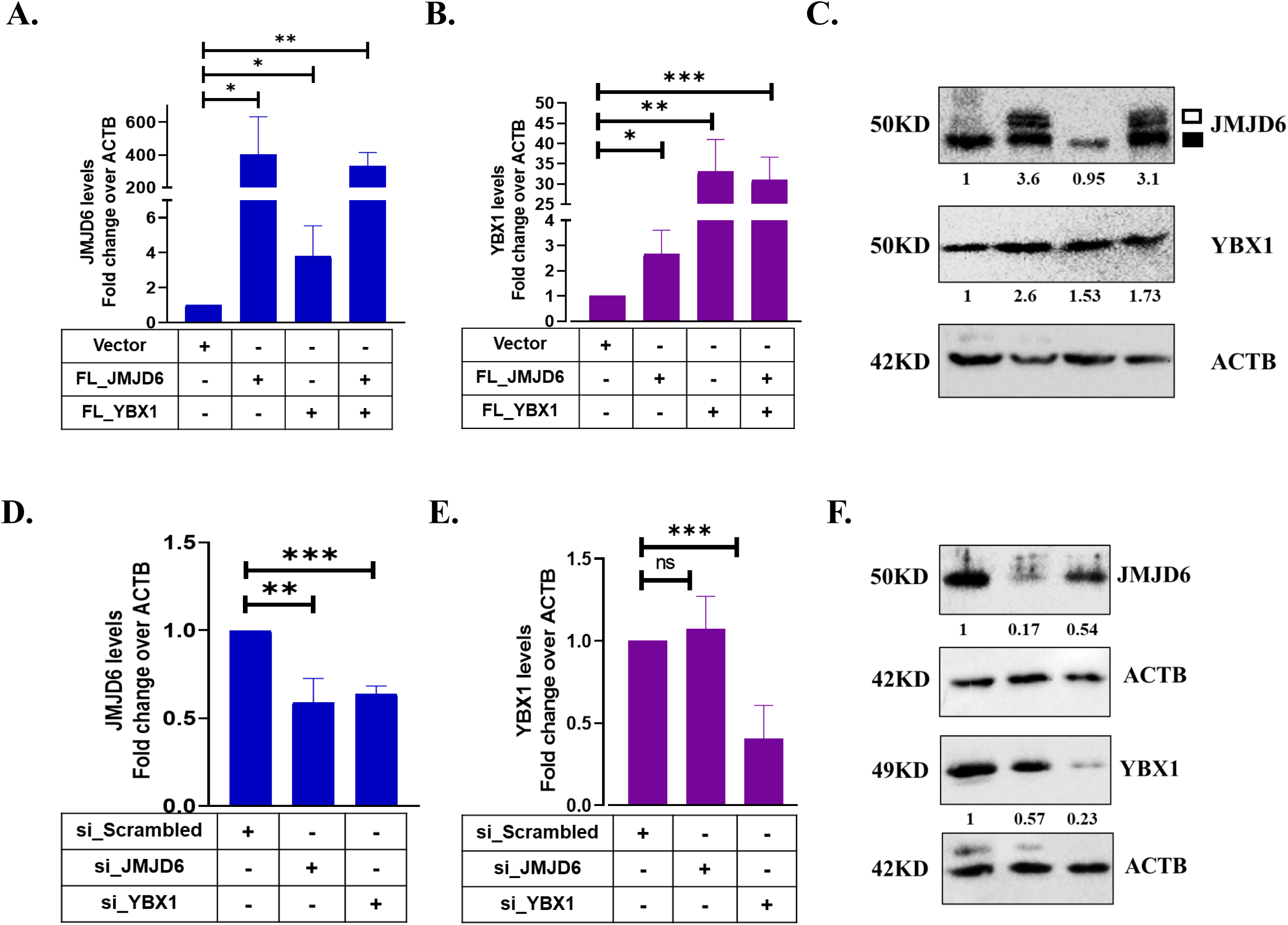
A) and B) Expression of JMJD6 and YBX1 following transient overexpression in MCF7. RNA levels in empty vector transfected cells are normalized to 1. C) Western blots for both proteins. Endogenous JMJD6 is marked by filled squares and empty squares indicate V5-tagged proteins (6). Numbers represent densitometric scanning data; D) and E) Expression of JMJD6 and YBX1 following siRNA mediated knock down in MCF7 cells. Expression in scrambled siRNA treated cells is normalized to 1. F) Western blots for both proteins. Numbers represent densitometric scanning data.

### JMJD6 and YBX1 interact with each other

To validate the mass spectrometry data, we first recombinantly co-expressed V5-tagged JMJD6 and Myc-tagged-YBX1 in HEK293 cells and performed Co-IP. Figure 3A shows that the exogenously expressed proteins interacted with one another (Figure 3A). Next, endogenously expressed proteins were subjected to Co-IP assays in MDA MB 231 and JOE cells. Results showed that both proteins interact with each other (Figure 3B-C). Since JMJD6 was nuclear in JOE cells and cytoplasmic in MDA MB 231 cells, and YBX1 was mostly localized in the cytoplasm but shuttles in and out of the nucleus, cell fractionation was undertaken. Co-IP using nuclear and cytoplasmic extracts across these cell lines showed that wherever the two proteins co-localized, they interacted with one another (Figure 3D, Supplementary figure 2A). Firstly, since both JMJD6 and YBX1 are known to interact with RNA and could interact via this intermediate, the co-IP experiments were conducted in presence and absence of RNAse in HEK293 cells. No difference was obtained in the two co-IPs suggesting that RNA presence may not be essential for these proteins to interact (Figure 3E). Further, JMJD6 and YBX1 protein were synthesized using cell free reticulocyte lysate systems (Supplementary figure 2b) and co-IP was carried out. Figure 3F shows positive interaction of these *in vitro* synthesized proteins (Figure 3F). It is highly likely both the proteins interact directly with each other.

**Figure 3:**
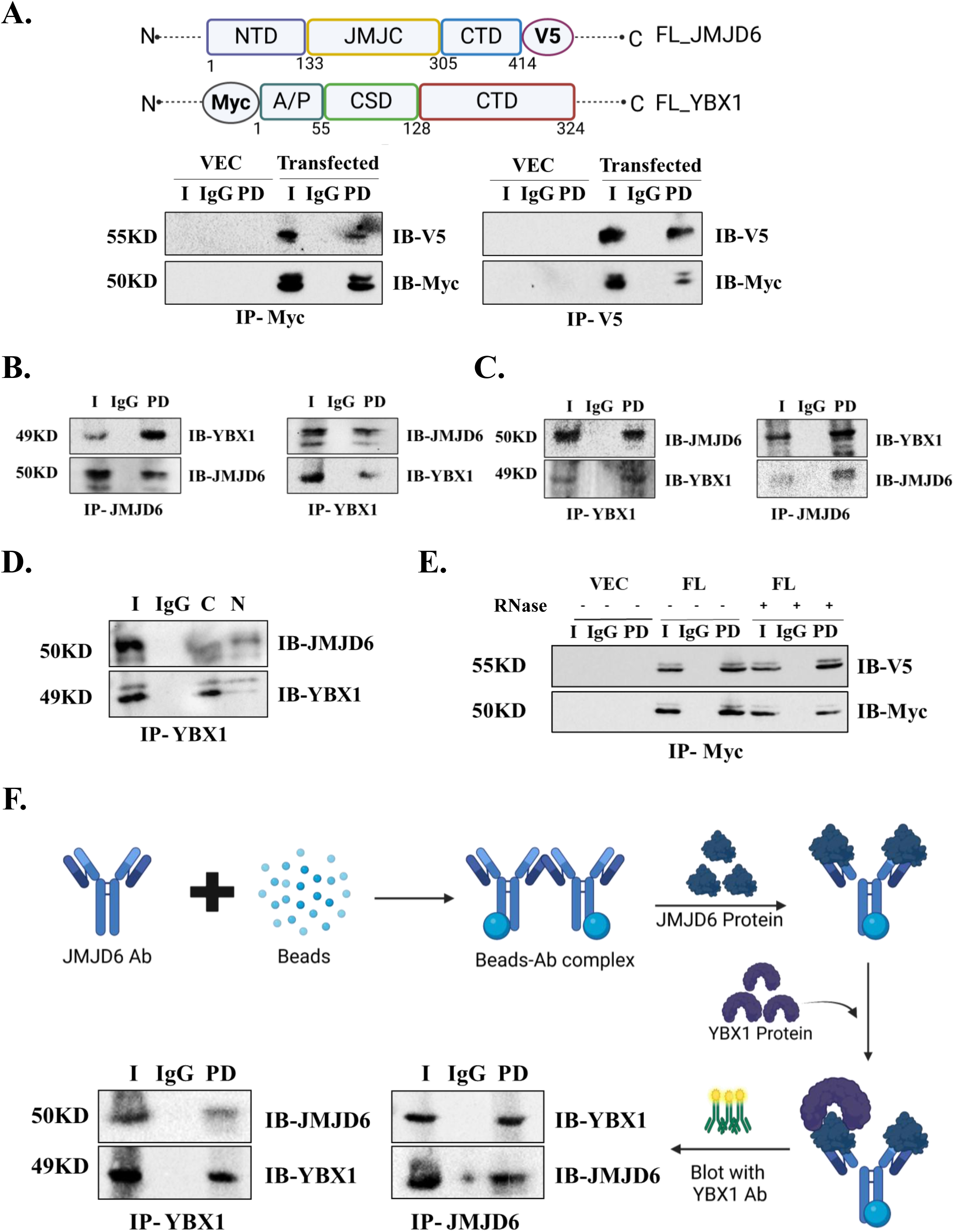
Interaction of JMJD6 and YBX1 A) CoIP using exogenously overexpressed proteins in HEK293 cell line; B) and C) CoIP with endogenous proteins in JOE and MDA MB 231 cells respectively, D) CoIP in nuclear-cytoplasmic extracts of JOE cells, E) CoIP using recombinantly expressed proteins with/without RNase treatment, F) Experimental design and CoIP with *in vitro* synthesized proteins (‘I’ represents Input, ‘PD’ denotes immune precipitated protein, ‘IB’= Antibody used for immunoblot, ‘IP’= Antibody used for pulldown

### Domain mapping of JMJD6 and YBX1

Next, the region of interaction between JMJD6 and YBX1 was mapped. Three deletion constructs of YBX1 with N-terminal Myc tag were constructed as follows-i) pYCTD (128-324 amino acid), ii) pYCSD-CTD (55-324 amino acid), iii) pYA/P-CSD (1-133 amino acid) (Figure 4A, supplementary figure 3A). We recombinantly expressed each of them with full-length JMJD6-V5 in HEK293 cells. Only pYA/P-CSD construct showed a positive interaction with full length JMJD6 (Figure 4B-D). We made two deletion constructs of JMJD6 namely pJΔCTD (1-305 amino acid) and pJMJC (J_133-305 amino acid), both with C-terminal V5 tag (Figure 4E, Supplementary figure 3B). We co-expressed them with full length YBX1-Myc tag. Upon immunoprecipitation with Myc, both JMJD6 constructs were detected by immunoblotting with V5 antibody. This denotes that JMJC domain of JMJD6 is sufficient to bind with YBX1 (Figure 4F-G). In conclusion, the A/P domain of the YBX1 was essential for interacting with JMJC domain of JMJD6.

**Figure 4:**
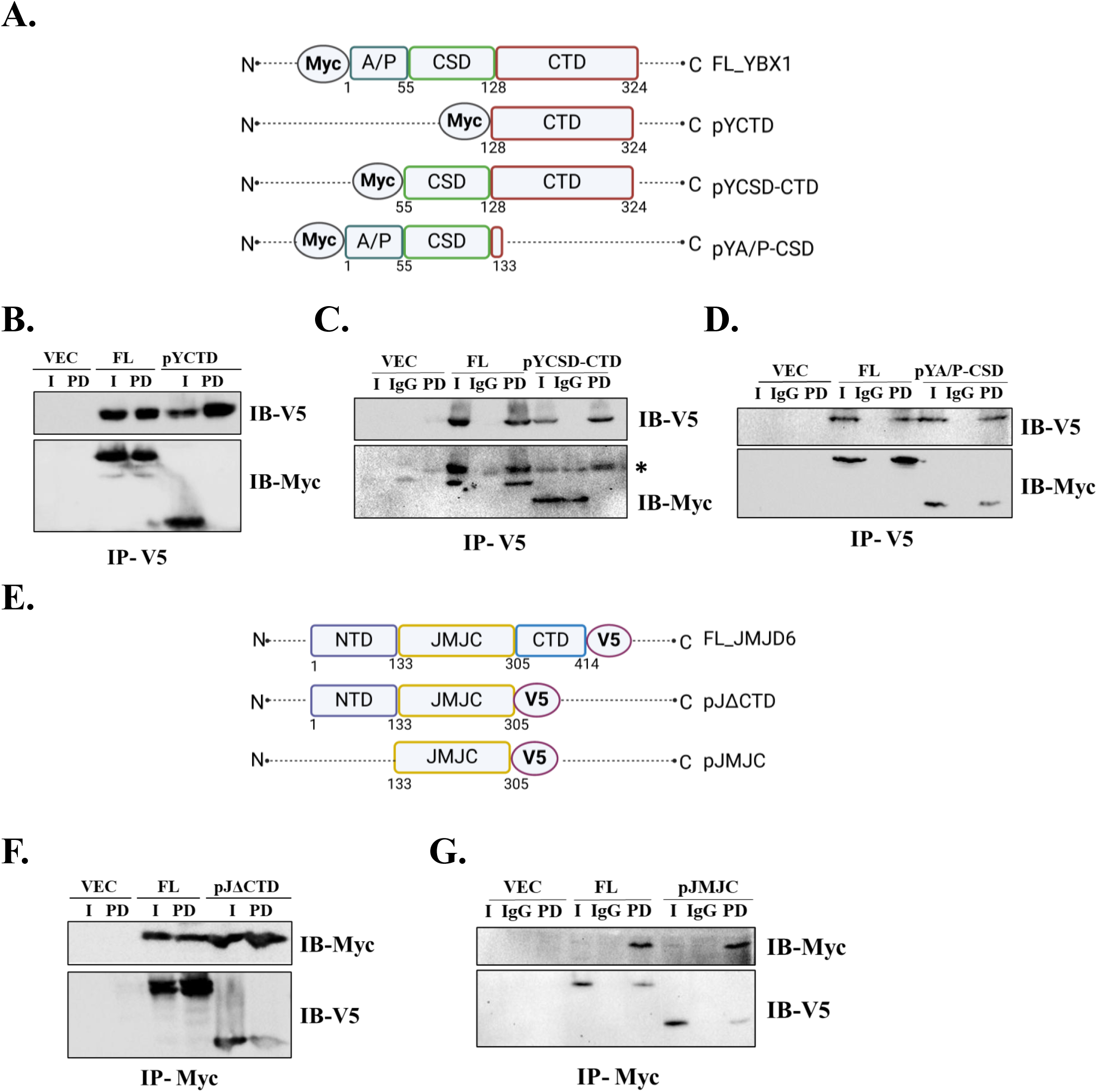
Domain mapping A) Myc-tagged deletion constructs of YBX1. CoIP of full length JMJD6 with B) pYCTD, C) pYCSD-CTD, D) pYA/P-CSD; E) Deletion constructs of JMJD6 tagged with V5. CoIP of full length YBX1 with F) pJΔCTD, and G) pJMJC (* denotes endogenous Myc)

### YBX1 binding induces the HOTAIR promoter

Since the two proteins interact with each other, we used two promoter constructs pHP216 and pHP123 to determine the effect of YBX1 on their activity (Figure 5A). Promoter assays were conducted in MCF7 cells after CRISPR-Cas9 mediated transient knock out of JMJD6 (JKO) and YBX1 (YKO) using pooled clones (Figure 5B, upper panel). Repeatedly, single clones of knock out cells were not obtained due to limited growth or death of KO cells. Therefore, decrease in protein expression was used to confirm partial KOs by western blots prior to promoter assays (Figure 5B, lower panel). A 50-70% reduction in luc activity was evident when either of the proteins was depleted as compared to the parental MCF7 cells. Since HOTAIR had enhanced activity in JOE cells, the effect of YBX1 KO and its effect on promoter constructs was checked in JOE and Vec cells. Strikingly, YBX1 affected both the basal and JMJD6 induced HOTAIR promoter activity (Figure 5C). However, as the YBX1 site was predicted in the (−216bp to -123bp) region, we expected that YBX1 would likely decrease the activity of only the pHP216 construct and not pHP123. These data first suggested that YBX1 may be instrumental in recruiting JMJD6 to its site and YBX1 was a more impactful regulator of the HOTAIR.

**Figure 5:**
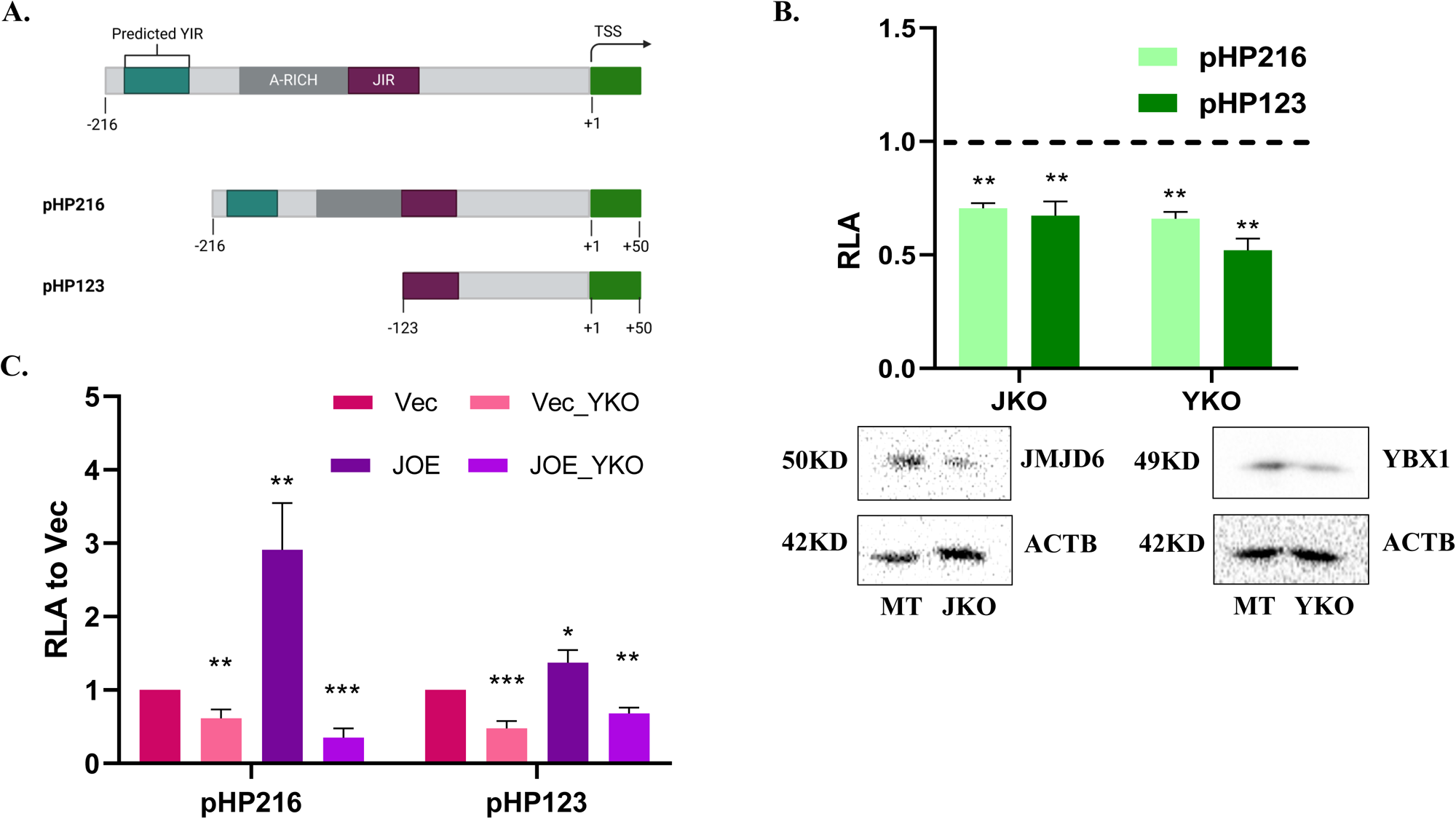
HOTAIR promoter regulation A) Details of HOTAIR promoter constructs; B) Luciferase assays in MCF7 cells with transient knockout of JMJD6 (JKO) and YBX1(YKO). Dotted line represents luciferase activity in empty vector transfected cells normalized to 1. Extent of protein depletion is shown by western blots in the lower panel; C) Luciferase assay in Vec and JOE cells after YBX1 KO.

### YBX1 interacts with HOTAIR promoter

Since YBX1 affected HOTAIR promoter activity, YBX1 ChIP was performed and the resultant DNA material was amplified with primers encompassing the 216 bp promoter proximal region of HOTAIR. YBX1 ChIP was validated using reported positive binding sites in the SOX2A and MET1 regulatory regions (Supplementary figure 3)(23,24). A substantial positive enrichment for YBX1 was observed in the proximal promoter region of HOTAIR (Figure 6A). Since the neighbouring region was identified by us as JIR, and both JMJD6 and YBX1 interact, we performed ChIP-re-ChIP assays to assess if both proteins simultaneously occupy this region. We pulled down chromatin using YBX1 antibody followed by sequential JMJD6 pull down (seqJMJD6) and vice versa (Figure 6B and C). PCR data showed significant enrichment using ChIP and re-ChIP material in both cases. Both proteins, therefore, are likely occupying nearby regions in the promoter region of HOTAIR. To determine if JMJD6 binding and/or activity was influenced by YBX1 binding close by, we performed YKO in JOE cells and chromatin from these cells was subjected to JMJD6 ChIP. (Figure 6D). The level of enrichment in YKO cells decreased substantially as compared to mock transfected JOE cells (MT). These data recapitulate the data of the luc assays and reinforce the idea that YBX1 helps in JMJD6 recruitment at the JIR site.

**Figure 6.**
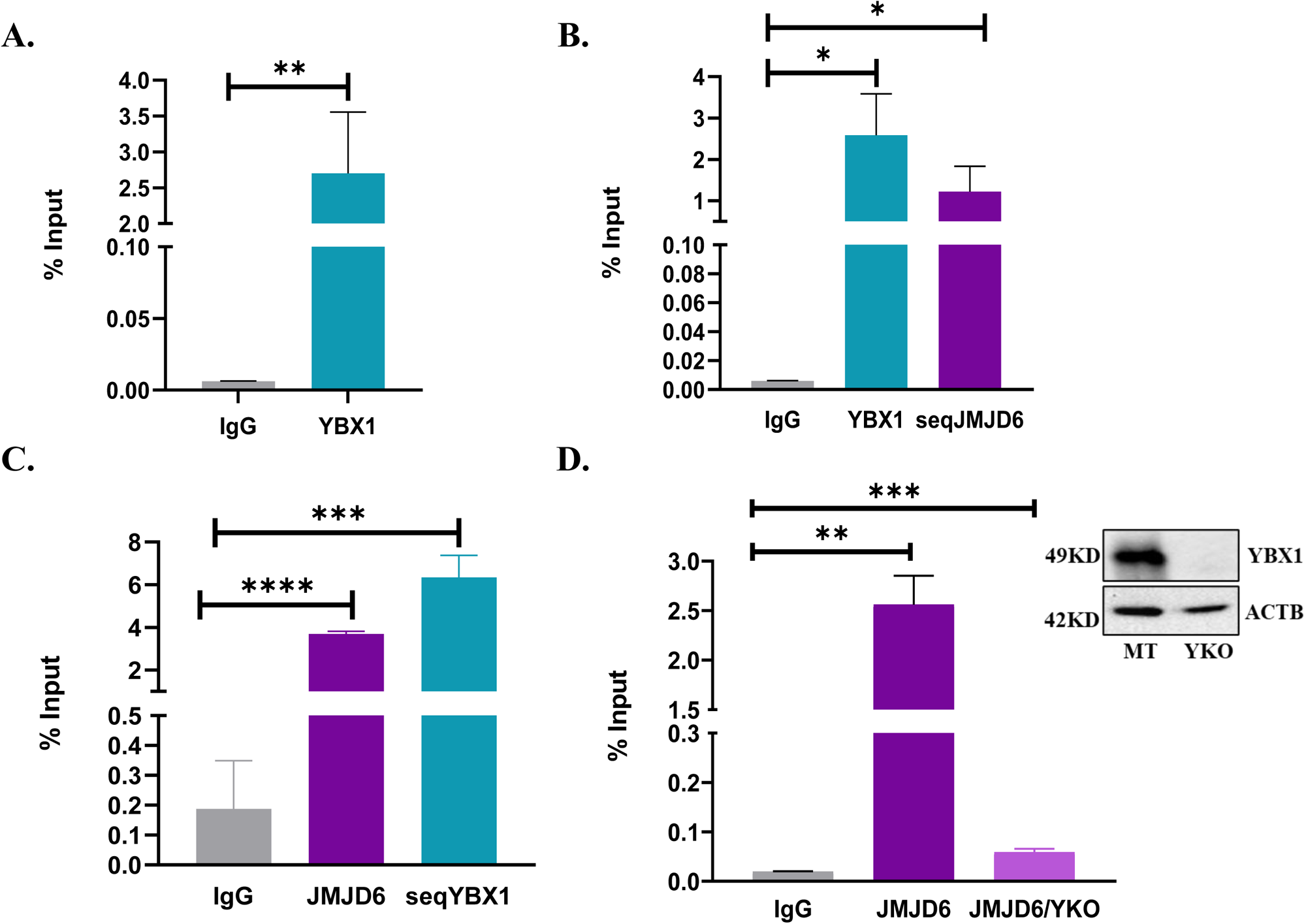
ChIP followed by sequential ChIP assays for HOTAIR promoter (−216 - +50 bp) A) YBX1 ChIP. B) YBX1 ChIP followed by JMJD6 pull down (seqJMJD6) and C) vice versa. D) JMJD6 ChIP in mock transfected (MT) and YKO cells. YBX1 protein levels are shown by western blot (inset).

### YBX1 interaction region

The potential site of YBX1 interaction (YIR) was between (−216 to -123 bp). We constructed two probes, one spanning this YIR and another to the known JIR (Supplementary table 1). Electromobility shift assays were performed using nuclear extracts from JOE cells. For potential YIR, two complexes appeared (Complex I and II) and the 50X excess of cold probe removed Complex II and reduced the intensity of Complex I (Figure 7A, left panel). Nuclear extracts from JOE_YKO cells completely abolished all complexes indicating that YBX1 was responsible for these shifts. As shown earlier, JIR region retarded the cognate probe (Complex III), 50X cold excess probe competed it out and loss of YBX1 resulted in significantly weakening the intensity of the retarded complex. Appearance of Complex IV was observed in the YKO lane, however, binding partner for this shift is unknown (Figure 7A, right panel). Immunoblot for JOE MT and YKO cells is shown in Figure 7B. These data conclude that YBX1 interacts with the potential YIR and indicates that JMJD6 binding to JIR is probably enhanced in the presence of YBX1.

**Figure 7:**
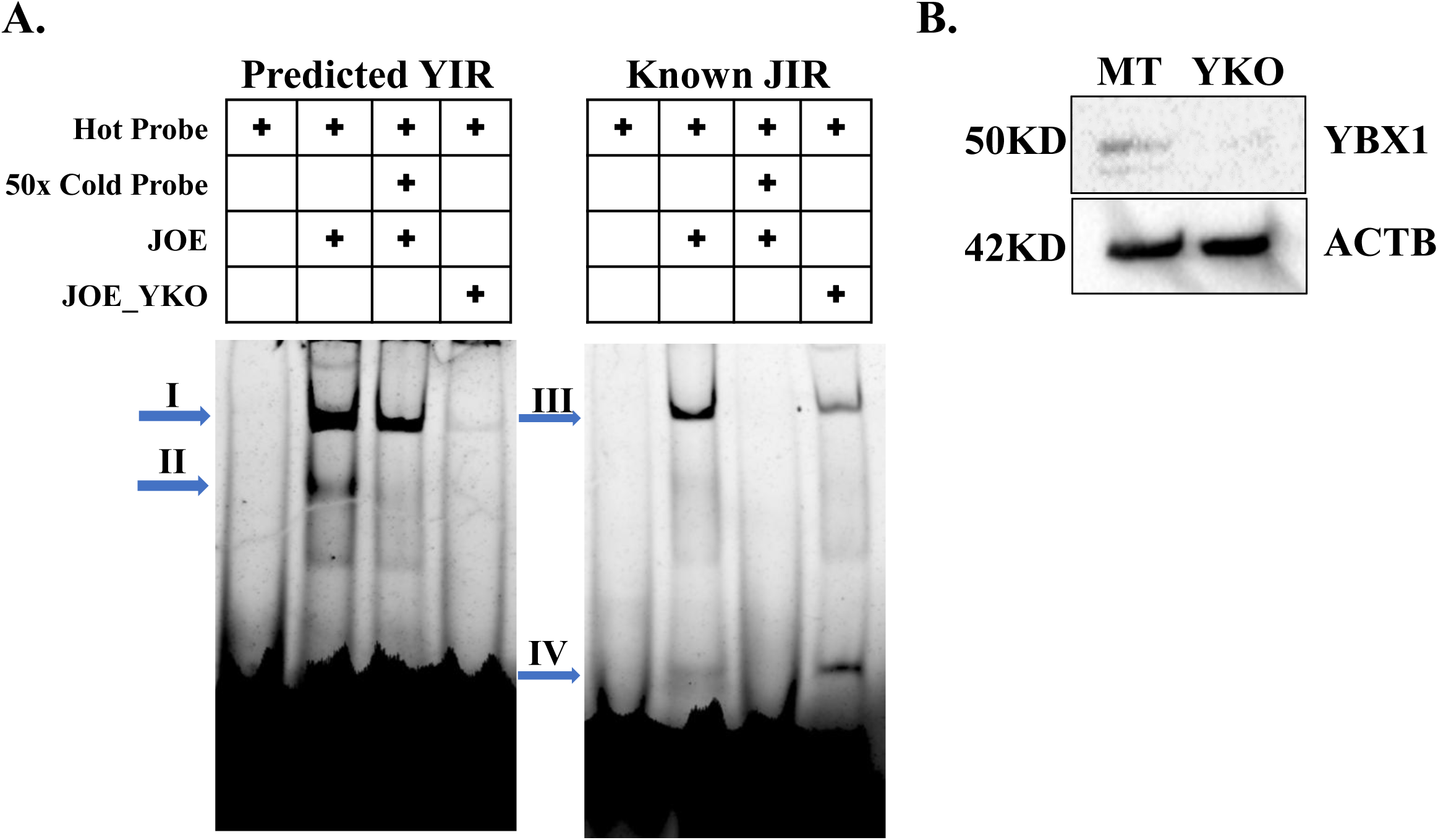
YBX1 interacting region of HOTAIR promoter A) EMSA of potential YIR and known JIR using nuclear extracts from JOE cells with and without competing cold probe and in the presence/absence of YBX1. Right-ward arrows denote complexes (I-IV). B) Western Blot to confirm depletion of YBX1 in JOE cells. (MT= Mock transfected cells)

### YBX1 positively regulates HOTAIR RNA expression

Quantitative real-time PCRs under various conditions of presence/absence of the two proteins were conducted to test if their binding results in change in HOTAIR expression. Depletion of either protein resulted in decreased HOTAIR RNA expression in MCF7 cells (Figure 8A). Transient over expression of JMJD6 and YBX1 increased the level of HOTAIR RNA in MCF7 cells (Figure 8B). Similarly, siRNA mediated depletion of YBX1 in JOE and Vec cells significantly reduced HOTAIR expression in both cells (Figure 8C, upper panel). Therefore, YBX1 affects the basal levels of HOTAIR RNA and high JMJD6 alone is not sufficient to maintain the elevated HOTAIR levels. This observation further supports YBX1 mediated recruitment of JMJD6. Also, it was observed from publicly available data that high expression of all 3 genes in breast cancer was more prone to high hazard ratio and poor survival in clinic (Figure 8D).

**Figure 8:**
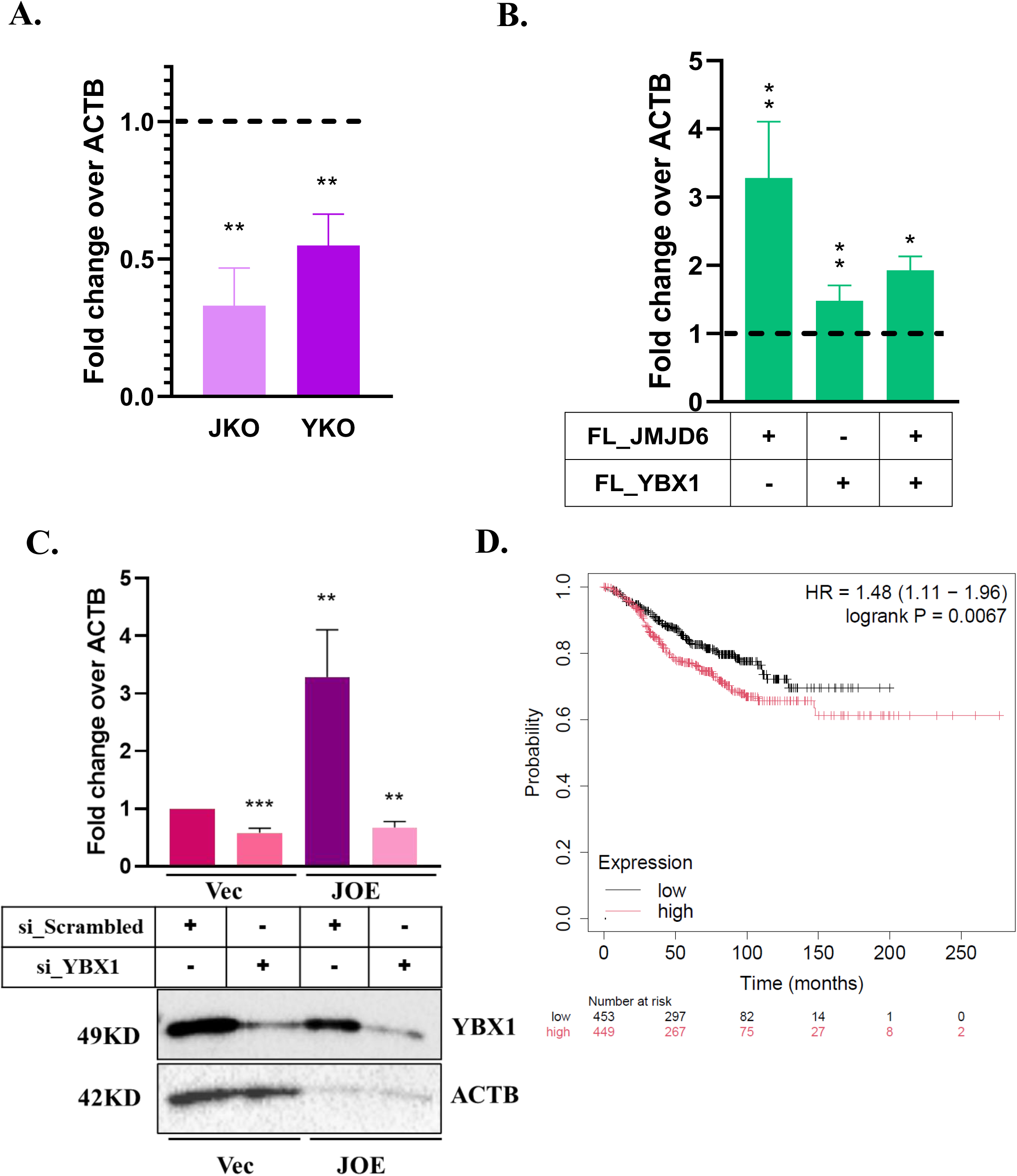
HOTAIR RNA expression A) qRT-PCR of HOTAIR in JKO and YKO cells. Dotted line represents HOTAIR levels in control mock transfected cells, normalised to 1. B) qRT-PCR of HOTAIR in MCF7 cells after transient overexpression of the two proteins. Expression of empty vector is marked as dotted line and converted to 1. C) qRT-PCR of HOTAIR in Vec and JOE cells following YBX1 siRNA mediated KD. si_Scrambled was used as control and converted to 1. Western blots are shown in lower panel. D) Survival analysis using all three genes (JMJD6, YBX1, HOTAIR), red line denotes high expression of all three and black line denotes low expression of all three genes. Lower panel represents patient number.

## Discussion

Earlier we identified a JMJD6 interacting region within 120 bases of the HOTAIR TSS which increased the basal promoter activity in the presence of JMJD6. Addition of another 130 bases upstream (−216 to +50 region) doubled JMJD6 induced luc activity. However, we were unable to find the factor responsible for this improvement. TRANSFAC predicted several proteins as potential interactors, but mutation of such regions failed to decrease the promoter activity (6). In the current study, we revisited this problem and utilized experimental ChIP data generated for 160 transcription factors in MCF7 by ENCODE. Only 4 factors displayed low enrichment in this region. Of these, our previous mutations encompassed EP300, ZBTB11 and FOXA1. The factor that remained unexplored was YBX1. YBX1 has potential for binding to DNA as well as RNA and it is known to regulate both gene transcription and translation (25,26). The ENCODE data tracks for YBX1 spanning the HOTAIR proximal promoter region were explored in the IGV browser (data not shown), and a potential YBX1 binding region was evident near the (−216 to -100 bp) region. This was approximately 60 bp upstream of JIR. In addition, previous studies identifying JMJD6 interacting proteins using JMJD6 IP listed YBX1 as one of the interacting partners (4,7). These two observations prompted us to hypothesize that YBX1 physically interacts with JMJD6 and regulates the HOTAIR promoter by binding near JIR.

Immunoblotting showed ample expression of YBX1in MCF7, Vec, JOE and MDA MB 231 cells. Surprisingly, although JMJD6 is a well characterised nuclear protein, MDA MB 231 cells had higher expression of this protein in the cytoplasm and this has been described earlier (21,22). YBX1 was present in both cytoplasm as well as nucleus in all cells tested. Considering the premise of the current work of regulation of HOTAIR promoter, we used Vec and JOE cells for promoter assays due to observed nuclear expression of both proteins but examined molecular interactions between them in multiple cells.

We tested if recombinant expression of JMJD6 in MCF7 cells (JOE cells) altered YBX1 RNA and protein levels. Both western blots and RT-PCR confirm JMJD6-mediated positive regulation of YBX1 in JOE and JMJD6 siRNA treated cells. Interestingly, downregulation of JMJD6 RNA and protein was evident following YBX1 siRNA treatment in MCF7 cells. In addition, transient over expression of both proteins showed enhancement of each other’s expression. Our manuscript does not explore the mechanisms involved in this regulation. However, both proteins are known to affect Myc levels in breast cancer cells and Myc upregulates JMJD6 by binding to its promoter (27). JMJD6 in turn induces MYC (11). YBX1 and Myc axis is known to negatively affect cancer cell apoptosis and YBX1 mediated stabilization of myc protein has been demonstrated (28). This data suggests that a reciprocal regulatory loop may be established between JMJD6 and YBX1 and MYC may serve a mechanistic role therein.

To validate the mass spectrometric data from JMJD6 IP, co-IPs were performed using JMJD6 and YBX1 antibodies followed by immunoblots to detect the interacting partner. Recombinantly overexpressed, *in vitro* synthesized and endogenous proteins consistently displayed positive protein-protein interactions. Further this interaction was also seen in MDA MB 231 cells where we found cytoplasmic localization of JMJD6. In Vec or MCF7 cells, this interaction could not be tested due to low expression of endogenous JMJD6 as compared to JOE and MDA MB 231 cells. Further, nuclear and cytoplasmic extracts in the latter cells showed protein-protein interaction in both cellular compartments. JMJD6 and YBX1 are both RNA binding proteins and to test if this interaction involves RNA, co-IP was performed in presence or absence of RNase. The binding of these two proteins was agnostic to the presence of RNase and treatment did not hamper their interaction. These experiments suggest that direct interaction between these proteins is highly possible.

Next, we looked for protein domains involved in the interaction between JMJD6 and YBX1. YBX1 has 3 major domains-alanine proline rich domain (A/P domain), cold shock domain (CSD), and C terminal domain (CTD). Deletion constructs pYCTD and pYCSD-CTD were unable to bind full length JMJD6. This suggests that binding is probably within the A/P domain. The A/P domain is about 55 amino acids and finite expression and IP could not be achieved using this construct. Another JMJC family member, JARID2, also interacts with YBX1 (29). This study successfully expressed A/P domain with 88 amino acids of CSD and similar clones were generated in this study (29). This construct, pYA/P-CSD, could bind JMJD6. This indicates A/P domain of YBX1 was required for JMJD6 interaction. Analysis of JMJD6 deletion constructs indicated that the JMJC domain could interact with pYA/P-CSD. We did not analyze any constructs of JMJD6 that were deleted for JMJD6 C-terminal domain since Cockman *et al* have shown that mass spectrometric analysis using CTD truncated JMJD6 (beyond amino acid number 345) did not affect YBX1 pull down (30). Overall, both studies suggest that the JMJC domain was sufficient for interaction with A/P domain of YBX1. Using Pepsite2 program, we scanned 30 amino acid peptides generated over this construct with +1 as the sliding window. The “QPPR” amino acid sequence of the A/P domain displayed low and highly negative ΔG values. Whether JMJD6 enzymatically modifies YBX1 via arginine demethylation of the QPPR sequence would be an interesting aspect to study. Previously, removal of arginine methylation by JMJD6 from ER and G3BP1 (a stress granule protein), enhanced their nuclear localization (9). More work in the future can be undertaken to explore if JMJD6 utilizes this strategy to concentrate YBX1 in the nucleus. This is important since several studies have shown that nuclear localization of YBX1 protein following phosphorylation by receptor tyrosine kinase effector proteins such as ERK1, promotes aggressive tumor behavior and cellular invasiveness. Similarly, binding with HOTAIR also enriches YBX1 nuclear amounts (18). While, JMJD6 is also a lysyl hydroxylase, but in a recent study by Cockman et al, YBX1 was not identified as an enzymatic target of JMJD6 (30).

After establishing the two proteins interact with each other, we tested if YBX1 had any role in HOTAIR promoter regulation using the pHP216 (−216 to +50 bp) and pHP123 (−123 to +50 bp) constructs. In MCF7 cells, transient KO of either JMJD6 or YBX1 resulted in about a 50-70 % loss of luc activity of both constructs. To estimate the contribution of YBX1 independently of JMJD6, we used Vec and JOE cells, since JMJD6 expression is marginal in MCF7, high in JOE but YBX1 is abundantly expressed in these cells. In both Vec_YKO and JOE_YKO cells, relative luciferase activity of pHP216 and pHP123 diminished. However, it was surprising that pHP123 luc activity decreased after YKO, since this construct lacked the potential YIR region present in pHP216. These observations along with the co-IP interaction data, gave a first clue that YBX1 may recruit JMJD6 to the HOTAIR promoter. While JMJD6 is capable of binding single stranded RNA, its ability to interact with DNA appears to be like a secondary binder, mediated via the DNA binding regions of interacting transcription factors (31).

Next, we studied if enrichment for the proximal promoter region could be obtained by YBX1 specific ChIP-PCR assay. Indeed, significant enrichment was observed. To ascertain if both factors co-occupied the promoter region simultaneously, sequential ChIP-re-ChIP assays were performed using chromatin material from JMJD6 ChIP for re-CHIP using YBX1 antibody and vice-versa. Both ChIP-re-ChIP assays indicated enrichment for both proteins suggesting they occupy neighbouring region(s) in the HOTAIR promoter. Interestingly, YKO resulted in decrease in JMJD6 occupancy showing dependency of JMJD6 on YBX1 for interacting with JIR. Next, we constructed two non-overlapping probes spanning the potential YIR and the known JIR. Gel-shift experiments identified 2 retarded complexes using nuclear extracts from JOE cells. Complex II could be competed out using 50X molar excess of cold probe whereas a slight decrease in signal, if at best, was observed for Complex I. Intriguingly, YKO in JOE cells completely abolished both complexes. This indicated that it was highly likely that YBX1 was at least one of the proteins that interacted with this region. More interesting were the complexes obtained when JIR probes were used in similar conditions. A single complex (III) was observed that was competed out by the cold probe. Surprisingly, YKO extracts decreased complex III formation, and another small complex (IV) was intensified. Appearance of such small complexes has been attributed to presence of histones in several other studies. Overall, these data confirm that YBX1 assisted JMJD6 binding to JIR and enhanced JMJD6 mediated HOTAIR promoter activity.

The question remained if the binding of YBX1 and JMJD6 resulted in increase in HOTAIR RNA expression. In several ways, we perturbed expression of both proteins and confirm that these proteins augment HOTAIR RNA expression. Particularly, in JOE cells, removal of YBX1 caused reduction of HOTAIR RNA indicating it participates in JMJD6 mediated upregulation. Interestingly, the YIR and JIR regions are separated by an A-rich region and we have previously shown that deletion of this region uncouples the requirement of JMJD6 for HOTAIR induction and results in very high activity of TATA-luc constructs. We and others have suggested that this could be explained by the interaction of repressor proteins such as IRF-1 with the A-rich motif (32). We found that in YKO cells, the expression of the repressor IRF-1 was highly induced (data not shown). Therefore, it is possible that YBX1 also enhances HOTAIR transcription independent of its binding by down modulating the expression of its repressor. Further studies are required to explore all the complexities involved in the regulation of HOTAIR.

Interestingly, HOTAIR lincRNA promoted phosphorylation of YBX1 and enhanced its nuclear localization. YBX1 is phosphorylated by several kinases involved in signal transduction pathways including ERK, AKT, p70S6 kinase and its nuclear localization is also dependent on the phase of cell cycle (33). Phosphorylation of YBX1 increased expression of genes involved in multidrug resistance (such as MDR1) leading to resistance to endocrine therapy drugs such as Tamoxifen. Similarly, high HOTAIR expression is also involved in drug resistance. We have previously shown that JOE cells are less sensitive to Tamoxifen and harbor high levels of phospho-ERK and receptor tyrosine kinase RET levels. Both contribute to drug resistance (34). YBX1 was expressed in in both nucleus and cytoplasm of MCF7 cells. It is plausible that JMJD6 induction of receptor tyrosine kinase RET and ERK1 contributes to enhanced YBX1 phosphorylation and translocation to the nucleus, in turn inducing HOTAIR transcription. Additionally, these data have shown that JMJD6 and YBX1 induce each other and JMJD6 recruitment is dependent on YBX1 presence. Based on these data, we propose that JMJD-YBX1-HOTAIR are engaged in a feed forward positive loop. Once established, these 3 genes may lead to aggressive tumor behaviour, poor response to therapy and ultimately poorer survival outcome in ER+ patients. JMJD6 inhibitor has been used to regress tumors in preclinical mouse models and several efforts are being made to design YBX1 inhibitors since it promotes breast cancer metastasis (35,36). Therefore, targeting either of these two proteins may be a helpful strategy to combat invasive and aggressive breast cancer.

## Declarations

Ethics Approval and Consent to participate-Human/Animals samples were not used in this study

## Availability of data and materials

NA

## Conflict of interest

The authors declare no conflict of interest

## Funding

This project is funded by the DST-SERB Power grant (SPG/2021/004755). AG is supported by PhD fellowship from DST-INSPIRE (IF180919). SB is supported by DST-SERB Power grant (SPG/2021/004755).

## Authors contributions

A Gupta: Data generation, visualization, analysis. S Bhardwaj: Data generation, visualization and curation. KV Desai: Conceptualization, Supervision, Project coordination, data generation, analysis. All authors contributed to writing of first draft, review and editing of final draft of manuscript.

## Acknowledgement

NIBMG genomics core facility is acknowledged for extending use of their instrumental facility.

## Notes

### Competing Interest Statement

The authors have declared no competing interest.

### Summary of Updates

1) Change in figure resolution to improve size 2) Correction for mislabeling in Figure 4

