## Supplementary figures and images for "JMJD6 and YBX1 physically interact and regulate HOTAIR proximal promoter"

### Supplementary figure 1

**Supplementary Figure 1**

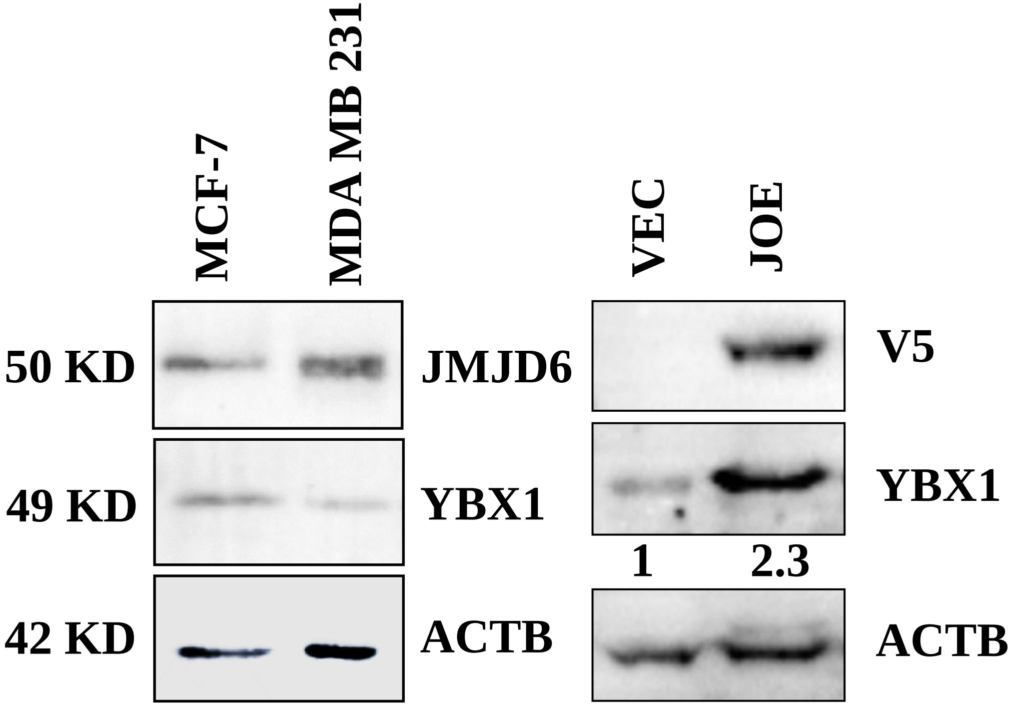

### Supplementary figure 2

##### Supplementary Figure 2

A.

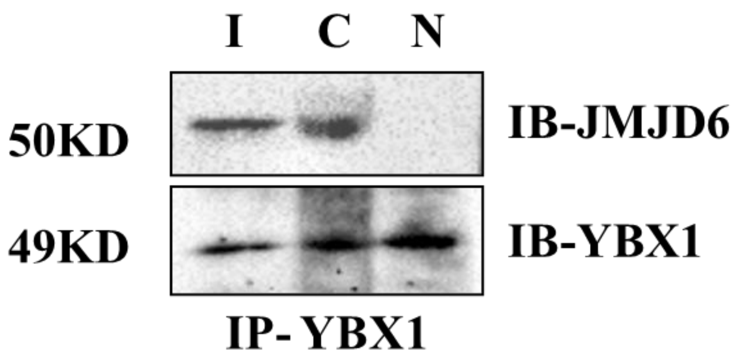


B.

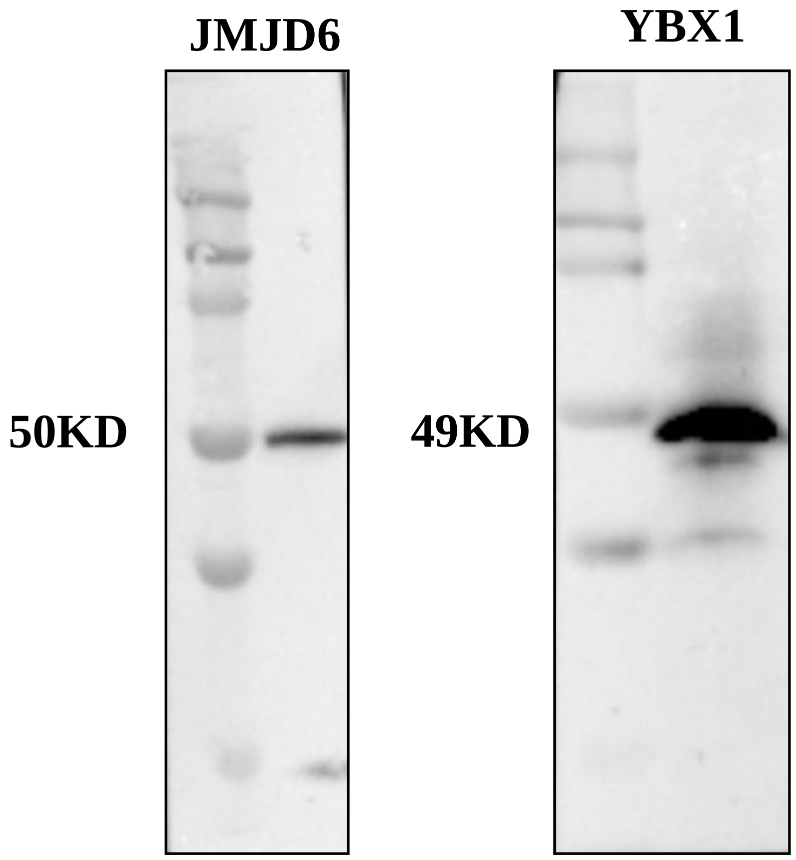

### Supplementary figure 3

##### Supplementary Figure 3

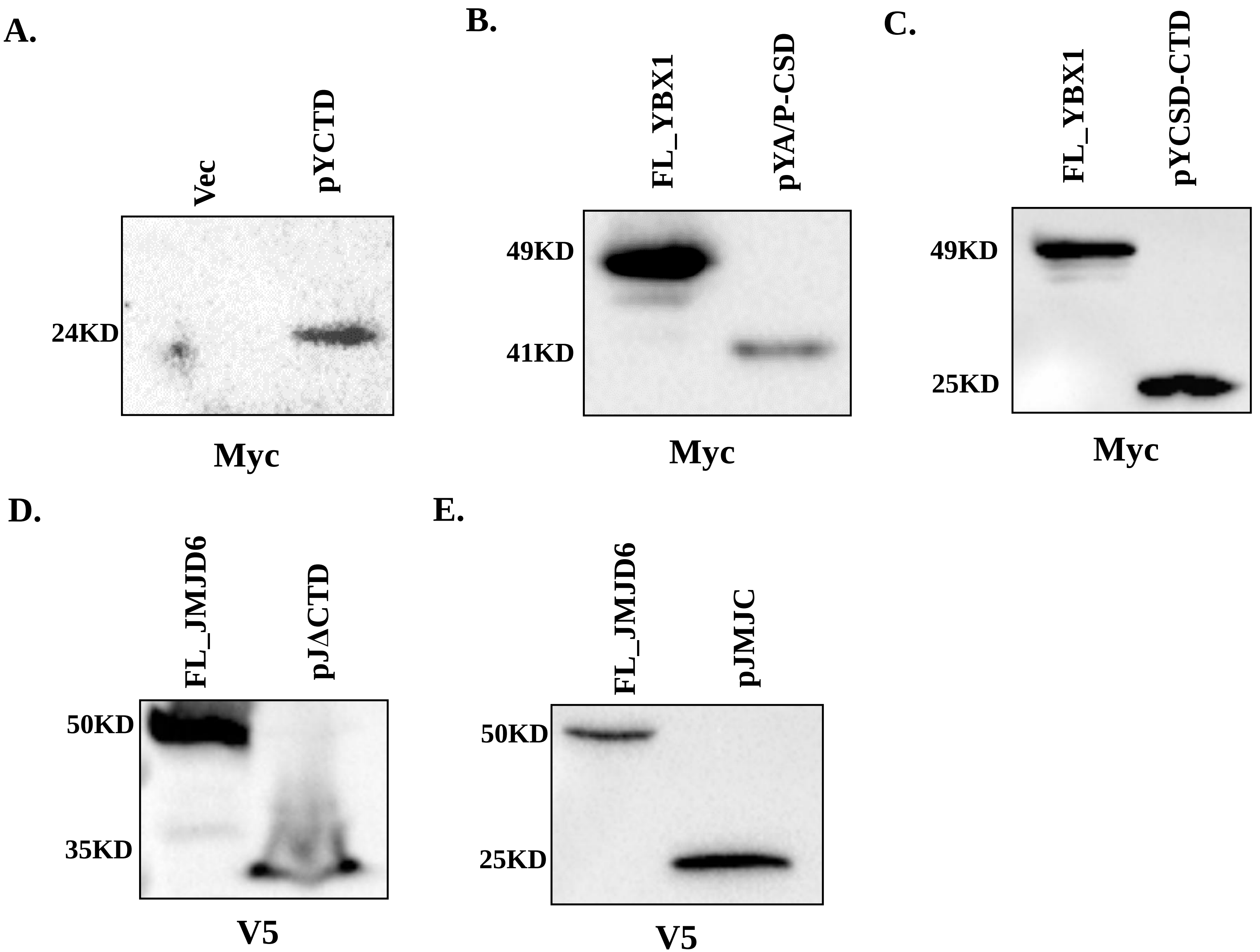

### Supplementary figure 4

#### Supplementary Figure 4

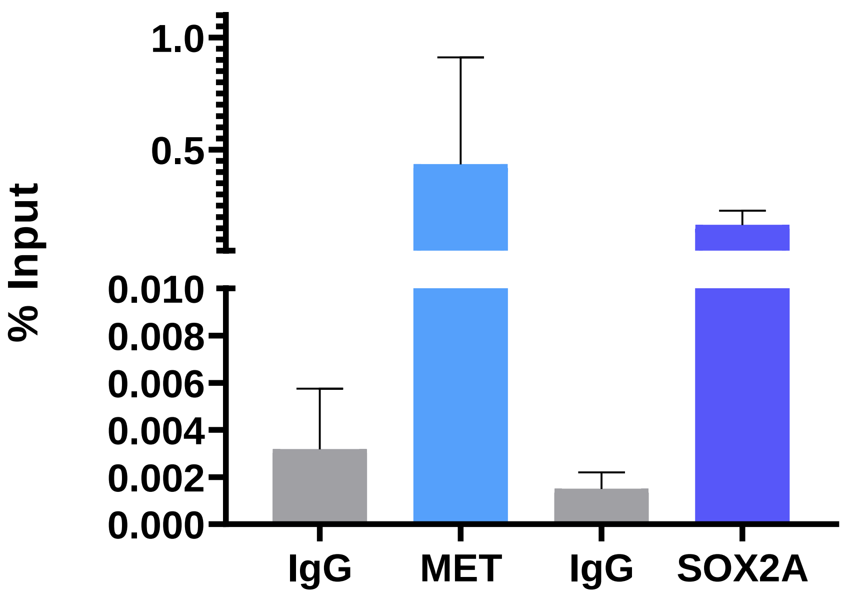
