## Supplementary table 1 for "JMJD6 and YBX1 physically interact and regulate HOTAIR proximal promoter"

| **Guide RNA sequences** | **Sequence (5'-3')** |
| --- | --- |
| JMJD6_sg_RNA | ACCGTCTTGTGCCATACCAC |
| YBX1_sg_RNA | GTGTACAAATACATCTTCCT |
| **qRT-PCR primer sequence** |  |
| JMJD6-RT-F | GGT TGA CCT TCA GGA GTC CAC |
| JMJD6-RT-R | TGC GCT CTT TGC TGA CAC AGT C |
| YBX1-RT-F | GCAGGAGAACAAGGTAGACCAG |
| YBX1-RT-R | CTTCATTGCCGTCCTCTCTAGG |
| Beta-actin-F | GAG CAC AGA GCC TCG CCT TT |
| Beta-actin-R | TCA TCA TCC ATG GTG AGC TGG |
| HOTAIR-RT-F | GGTAGAAAAAGCAACCACGAAGC |
| HOTAIR-RT-R | ACATAAACCTCTGTCTGTGAGTGCC |
| **ChIP primer sequence** |  |
| HOTAIR-F-216 | CGAGCTCGAAAAGAGAGGGGTGGGAAGG |
| HOTAIR-R-50 | CCGCTCGAGAGTCCTCACTGTGGAAGCTTT |
| MET-1-F | TTGACCTTCACACACCCAGAT |
| MET-1-R | TTCTGAGTTTGAGTGCCATGA |
| SOX2A-F | GAGAGAAA AAGGAGAACCTTCG |
| SOX2A-R | ACGGTGCATTG TTTTGTTCC |
| **EMSA Probe sequence** |  |
| Cy-5 universal | Cy5-GTGCCCTGGTCTGG-Cy5 |
| Predicted YIR-F | GAAAAGAGAGGGGTGGGAAGGCATGGGGTGAA AAA |
| Predicted YIR-R | TTTTTCACCCCATGCCTTCCCACCCCTCTCT TTTCCCAGACCAGGGCAC |
| Known-JIR-F | CTTTTGCCCCCAGCAAGAATCATTTGT |
| Known-JIR-F | ACAAATGATTCTTGCTGGGGGCAAAAGCCA GACCAGGGCAC |
| **Primers for deletion constructs** |  |
| HIND3 MYC CSD F | CCCAAGCTTATGGAACAAAAACTCATCTCAGAAGAGGATCTGAAGAAGGTCATCGCA |
| HIND3 MYC CTD F | CCCAAGCTTATGGAACAAAAACTCATCTCAGAAGAGGATCTGGGTGTTCCAGTTCAA |
| XHO1 CTD R | CCGCTCGAGTTACTCAGCCCCGCCCTGCTC |
| J1F | GAGGTACCATGAACCACAAGAGC |
| J133F | GAGGTACCATGGACAGCAGCTATGGT |
| J305F | GAGGTACCATGCCAAAGTTATCAAGG |
| J330V5R1 | CGATATCCTACGTAGAATCGAGACCGAGGAG |
